# ReproTox-KG: Toxicology Knowledge Graph for Structural Birth Defects

**DOI:** 10.1101/2022.09.15.508198

**Authors:** John Erol Evangelista, Daniel J. B. Clarke, Zhuorui Xie, Giacomo B. Marino, Vivian Utti, Taha M. Ahooyi, Sherry L. Jenkins, Deanne Taylor, Cristian G. Bologa, Jeremy J. Yang, Jessica L. Binder, Praveen Kumar, Christophe G. Lambert, Jeffrey S. Grethe, Eric Wenger, Tudor I. Oprea, Bernard de Bono, Avi Ma’ayan

**Affiliations:** Department of Pharmacological Sciences, Mount Sinai Center for Bioinformatics, Icahn School of Medicine at Mount Sinai, New York, NY 10029 USA; The Children’s Hospital of Philadelphia, Department of Biomedical and Health Informatics; Department of Pediatrics, University of Pennsylvania Perelman School of Medicine, Philadelphia, PA 19104, USA; Department of Internal Medicine, Division of Translational Informatics, University of New Mexico, Albuquerque, NM 87131; Department of Medicine, University of California San Diego, La Jolla, CA 92093, USA; Auckland Bioengineering Institute, University of Auckland, Auckland, New Zealand; Roivant Sciences Inc, Boston, MA 02210, USA

## Abstract

Birth defects are functional and structural abnormalities that impact 1 in 33 births in the United States. Birth defects have been attributed to genetic as well as other factors, but for most birth defects there are no known causes. Small molecule drugs, cosmetics, foods, and environmental pollutants may cause birth defects when the mother is exposed to them during pregnancy. These molecules may interfere with the process of normal fetal development. To characterize associations between small molecule compounds and their potential to induce specific birth abnormalities, we gathered knowledge from multiple sources to construct a reproductive toxicity Knowledge Graph (ReproTox-KG) with an initial focus on associations between birth defects, drugs, and genes. Specifically, to construct ReproTox-KG we gathered data from drug/birth-defect associations from co-mentions in published abstracts, gene/birth-defect associations from genetic studies, drug- and preclinical-compound-induced gene expression data, known drug targets, genetic burden scores for all human genes, and placental crossing scores for all small molecules in ReproTox-KG. Using the data stored within ReproTox-KG, we scored 30,000 preclinical small molecules for their potential to induce birth defects. Querying the ReproTox-KG, we identified over 500 birth-defect/gene/drug cliques that can be used to explain molecular mechanisms for drug-induced birth defects. The ReproTox-KG is provided as curated tables and via a web-based user interface that can enable users to explore the associations between birth defects, approved and preclinical drugs, and human genes.

## Introduction

The United States Department of Labor’s Occupational Safety and Health Administration [1] defines reproductive toxicity as a characteristic of “substances or agents that may affect the reproductive health of women or men, or the ability of couples to have healthy children. These hazards may cause problems such as infertility, miscarriage, and birth defects.” The prevention and clinical management of reproductive toxicity caused by chemical agents [2] requires the combined expertise from several medical fields including: public health and occupational health to protect against environmental/occupational toxins that lead to miscarriage [3], food and drug regulatory medicine to avoid drug teratogenicity or toxins in food that impact fertility, as well as clinical genetics, obstetrics, gynecology, and pediatrics to screen, prevent, monitor, and manage birth defects. This multidisciplinary nature of reproductive health is challenging. For instance, prescribing drugs in pregnancy remains a complex and controversial issue for mothers and physicians [4]. A key challenge to prescribing for the gravid patient is that recommendations are based on limited human pharmacological data and conflicting cases of adverse outcomes, given that pregnant populations are routinely excluded from randomized controlled trials [5]. Multidisciplinary challenges and data availability limitations are also key considerations in the prediction of drug-drug interactions [6,7] that may impact reproductive health [8,9]. As a result, birth defects, which in the US account for 3% of births [10] and 20% of infant deaths [11,12] are still mostly poorly understood.

In recent years, knowledge graphs have gained popularity as a productive approach to integrate data from multiple sources to organize information and gain new knowledge [13]. Knowledge graph databases store information about the semantic relationships between objects and represent relationships and events as triples: subject->predicate->object, for example, chicken->lays->eggs. Once these assertions are combined, they form a network made of nodes and edges and this establishes the knowledge graph. Once data from multiple sources are organized in a knowledge graph, it can be queried to extract subgraphs that can illuminate unexpected associations between entities. Integrated data organized into knowledge graphs can be used as input into graph embedding algorithms [14] that aim at predicting missing and novel associations that are not present in the original knowledge graph. Such an approach is increasingly applied in the domain of drug discovery [15]. In this study, we aimed at combining knowledge about birth defects with knowledge about genes and drugs to identify potential molecular mechanisms for known birth defects and predict birth defects for preclinical drugs and other small molecules. Specifically, we assembled evidence about the functional relationships between birth defect phenotypes and gene products and small molecules that are implicated in their occurrence from multiple sources. We ranked genes based on their association with pathogenicity; predicted the likelihood of small molecules to cross the placental barrier, and induce birth defects, using unsupervised learning; assembled knowledge about known drug targets for marked drugs [16]; and abstracted knowledge about the effects of drugs and preclinical small molecules on gene expression [17]. All this data is serialized into a knowledge graph representation and provided for access via a user-friendly web-based user interface.

## Methods

### Curating phenotypic terms relevant to birth defects

Human abnormal morphology of the great vessels, heart, and central nervous system (CNS) phenotypes were obtained from the EMBL-EBI Ontology Lookup Service (OLS) human phenotype ontology (HPO) v2021-10-10 [18]. To this end we considered the parent terms HP:0030962 (Abnormal morphology of the great vessels), HP:0001627 (Abnormal heart morphology) and HP:0012639 (Abnormal nervous system morphology) and extracted all the child nodes. The phenotype terms were then filtered based on relevance to birth defects. In all, 166, 193 and 252 phenotype terms were retained for great vessels, heart, and CNS respectively. In addition, 36 major birth defect terms were extracted from the Center for Disease Control and Prevention (CDC) website [19] on January 6, 2022.

To enhance the consistent representation of the above phenotypic terms, and to link these birth defects with knowledge about the appropriate anatomical entities involved with these pathologies, we manually curated the HPO terms onto an anatomy connectivity knowledge graph. The schema adopted by this graph is based on the ApiNATOMY knowledge representation [20,21], which was developed as part of the SPARC [22] connectivity mapping effort. The ApiNATOMY subgraph within the ReproTox KG provides links to knowledge about constituent anatomical structures such as cell types that may be involved in the birth defect mechanisms, as well as representations of abnormal anatomical organizations that typify these pathological phenotypes.

### Curating small molecules associated with birth defects

Manually curated known teratogens and xenobiotics that cause birth defects were extracted from various sources including Google searches and PubMed MeSH terms. We also used DrugCentral [23], which is an online drug information resource to query FDA D and X category drugs and their associated Simplified Molecular Input Line Entry System (SMILES) with absorption, distribution, metabolism, excretion, and toxicity (ADMET) properties. FDA approved drugs classified as X or D are drugs with evidence of inducing birth defects in humans and animal models. X category drugs should not be taken during pregnancy, while category D drugs should be avoided as much as possible. SMILES compound representations and names of active pharmaceutical ingredients were retrieved. We then utilized DrugCentral again to query birth defect terms that are within the FDA Adverse Event Reporting System (FAERS) [24] and mapped associated drugs with a likelihood ratio (LLR) cutoff of LLR > 2*LLRT. In addition, using DrugShot [25], we queried each CDC birth defect term through PubMed to extract PMIDs associated with each birth defect term. Abstracts associated with these PubMed IDs were mined to extract drug PubChem IDs based on co-mentions of the birth defect with a drug. The 30 most frequently occurring drugs for each birth defect were retained as the drug sets for each birth defect.

### Evidence implicating genes with birth defects

Given the curated phenotype lists described above, human phenotype-gene associations were retrieved from multiple sources, including Online Mendelian Inheritance in Man (OMIM) [26], Orphanet [27], ClinVar [28], DISEASES [29], DatabasE of genomiC varIation and Phenotype in Humans using Ensembl Resources (DECIPHER) [30], the American Heart Association (AHA) [31], and Geneshot [32]. From OMIM and Orphanet human phenotype-gene associations were obtained from the Jackson Laboratory HPO database (hpo.jax.org, October 2021 release), providing curated links between HPO terms and human genes. The OMIM and Orphanet-based HPO-term gene associations were retrieved for the human abnormal morphology of the great vessels, heart, and CNS phenotypes. Gene-birth defect associations were also obtained from ClinVar human genetic variants-phenotype submission summary dataset (v2021-11-03) [28]. This dataset was utilized to extract relationships between human genes harboring a pathogenic variant and their associated phenotypes given the birth defect phenotypes described above. Only genes with pathogenic variants and variants affecting a single gene were considered: that is, variants affecting multiple genes were excluded, due to the complexities in interpreting the relationships between their affected subset of genes and associated human diseases. The ClinVar-based HPO-gene associations were compiled for the human abnormal morphology of the great vessels, heart, and CNS phenotypes. Literature-based human disease-gene associations were obtained from the DISEASES portal [29]. This dataset contains disease-gene associations text-mined from literature and genome-wide association studies. The disease ontology identifier (DOID) and ICD-10 codes listed in this database were converted to HPO terms discussed above, which were used to filter the gene-term associations. The DECIPHER [30] provided this study with a curated list of genes reported to be associated with developmental disorders, processed by expert clinicians as a part of the Deciphering Developmental Disorders (DDD) study [33] to facilitate clinical feedback of likely causal variants. The DECIPHER-based HPO-gene associations were compiled for the human abnormal morphology of the heart, and CNS phenotypes. We included a dataset of human congenital heart disease-associated genes associated with syndromic, non-syndromic, and ciliopathic cardiac disorders that was recently published by the AHA as general guidance for genetic testing by practitioners in 2018 [31]. Finally, using the Geneshot API [32], we queried each one of the 36 CDC birth defect terms through PubMed to extract PMIDs associated with each term. These PubMed IDs were converted into genes using the AutoRIF option. The most frequently occurring genes were retained as gene sets for each birth defect.

### Linking small molecule and drugs to their known targets

Drugs and small molecules that have known targets were extracted from the TCRD database [16] and converted into KG assertions. Only compounds with a defined structure were included because other substances do not have PubChem [34] chemical IDs. In addition, only human targets were included, and only single gene/protein targets were included excluding some multi-component ion channels and transporters. Properties such as SMILES, binding affinity, original source, PubChem IDs, and common names are provided for each drug.

### Linking small molecules to genes based on changes in gene expression

The ReproTox-KG holds knowledge about most FDA-approved drugs and over 30,000 preclinical small molecules profiled by the LINCS program for their effects on the transcriptome of selected human cell lines [17]. To extract a set of genes that are up- or down-regulated by each drug and small molecule profiled by the L1000 assay for LINCS, we computed the mean of the Characteristic Direction [35] gene expression vector for each drug in the LINCS L1000 chemical perturbation signature dataset downloaded from SigCom LINCS [36]. We then retained the top 25 up- and down-regulated genes for each drug.

### Drug-drug similarity based on gene expression and chemical structure

To enable drug-drug similarity search across the ReproTox-KG, and to perform the unsupervised machine learning predictions, we developed two drug-drug similarity matrices, one based on structure, and one based on gene expression similarity. The drug-drug similarity matrix based on gene expression vector similarity was computed by transforming the consensus signatures described above using cosine similarity, comparing all pairs of consensus drug gene-expression vectors to produce a square matrix where the value at *(i,j)* is the gene expression-based cosine similarity between the drugs at row *i* and column *j*. The matrices that contain the consensus signatures for all drugs and small molecules, and the drug-drug similarity matrix are available for download from SigCom LINCS [36] and the ReproTox-KG download page. To create drug-drug similarity based on chemical structure similarity, we first converted the SMILES strings of each compound to a binary feature vector using the Morgan fingerprint (2048 bits) method [37] with radius 2, 3, and 4 was implemented in RDKit [38]. Other chemical structure similarity methods such as MACCS, Avalon, Atom Pair, RDKit with maxPath 2 and 4, and Topological fingerprints using FingerprintMol were tested. Next, we computed the inverse document frequency (IDF) between all pairs of drug vectors and the distance measure between each pair of drugs. We found that IDF performs much better than the standard Tanimoto similarity measure typically used for quantifying the similarity between pairs of compounds (data not shown). The resultant matrix of drug-drug similarity based on chemical structure is available from the ReproTox-KG download page. Chemical structure-based similarity search was also implemented using a workflow which queries the KG for compounds and generates fingerprints and similarity measures at runtime, for additional flexibility and interoperability. The CFChemDb database and development system was employed, with source code available at https://github.com/unmtransinfo/CFChemDb.

### Gene intolerance scoring

Gene intolerance scores were introduced to the knowledge graph from three main sources as measures of gene essentiality in human health and disease. Such intolerance scores are based on large-scale human exome sequencing projects and reflect the order of magnitude of the impact of negative selections on human genes, as their associated variants are filtered out from the human population most probably due to lack of ability of carrying individuals to pass them down (reproductive disadvantages). In this sense, the observed frequency of such genetic variants and their deviation from their expected values are computed through different methods and captured via the intolerance scores. In this section we include some technical details on the ingested intolerance scores. Gene intolerance scores were calculated utilizing three scoring systems assessing three interrelated intolerance measures, namely, haplo-insufficiency, triplo-sensitivity (collectively regarded as dosage sensitivity), and general intolerance. Augmented with population-wide whole exome sequencing, each of these measures aims to quantify the magnitude of consequences of expected and observed mutations within a gene and present it as a single value assigned to that gene. Probability of being loss of function intolerant (pLI) scores [39] for 18,225 human genes were obtained from a large-scale study conducted by the Exome Aggregation Consortium (ExAC) [40] using a database of exomes from 60,706 unrelated individuals sequenced as part of various disease-specific and population genetic studies. The basis for calculating pLI is to account for the frequency of observing disease-associated isoforms of the protein encoded by the gene which is resulted from a variant in its coding region. Therefore, the absence or low frequency of such variants implies higher gene intolerance to mutations and higher pLI. According to this study, 3,230 genes lacked almost any predicted protein-truncating variants (PTVs), signifying the variant effect on the emergence of a reproductively disadvantageous phenotype. In this regard, the pLI scores are defined over the range [0,1] as an estimate of the probability that a candidate gene is intolerant to a deleterious mutation. Specifically, larger values of pLI (closer to 1) are correlated with higher intolerance of the gene to mutations with genes having pLI ≥ 0.9 considered as highly intolerant genes. It should be noted that the opposite case is not necessarily true; that is, genes with low pLI could also be associated with lethal phenotypes or higher likelihood of pathogenicity depending on the stage of the human lifespan in which the gene expression plays a key biological role. Residual Variant Intolerance Score (RVIS) [41] values of 16,956 human genes were adopted from a large-scale analysis that processed 6,503 human whole exome sequences made available by the NHLBI Exome Sequencing Project (ESP) [42]. The scoring system developed in this study aimed to assess the functional mutations in the genic regions compared to all neutral variations that could occur in a gene and accordingly rank genes as a measure of intolerance to loss of function mutations. As an attempt to prioritize genes based on their likelihood of influencing disease/abnormal phenotype, the RVIS measures the deviation between the observed and predicted functional variants considering the total number of common variations in the target genes. The resulting scores were then compared with the available information on whether the gene causes any known mendelian diseases. In this sense, genes with higher functional mutations to total variant sites ratio will be considered more tolerant and vice versa [41]. Dosage sensitivity scores [43] such as haploinsufficiency and triplo-sensitivity are commonly used as measures of gene intolerance and disease association caused by low-frequency high-impact deletions and duplications known as rare copy number variants (rCNVs). Estimated haploinsufficiency and triplo-sensitivity for 17,263 of human genes were presented by a recent large-scale study meta-analyzing 753,994 individuals with neurological disease phenotypes [43]. This database presents an unprecedented knowledge repository on gene intolerance to duplication, whereas haploinsufficiency provides information on whether a heterozygous variant could lead to insufficient expression of the respective protein and its subsequent pathogenicity. In this sense, haploinsufficiency is an autosomal dominant gene action. This study characterizes 3,006 haplo-insufficient and 295 triplo-sensitive genes. The provided scores can effectively be utilized in gene prioritization by their potential loss-of-function or gain-of-function through the introduction of de-novo rCNVs as opposed to the point mutations.

### Gene-gene similarity based on co-expression

Gene-gene similarity associations were obtained from the human gene-gene correlation matrix provided by the ARCHS4 resource [44]. The matrix stores the Pearson correlation coefficient between genes across bulk RNA-seq expression samples uniformly processed by the ARCHS4 pipeline. Genes were filtered to include only protein-coding genes to keep the size of the graph manageable, and for each of the 17,966 genes, the top five most positively and most negatively correlated genes based on the correlation coefficients were extracted for a total of 170,819 edges. Each edge was weighted by the correlation coefficient between the two connected genes. These gene-gene associations were then integrated into the ReproTox KG. From these associations, it may be possible to identify novel genes that are potentially affected by known teratogens and discover how the role that different groups of genes may play in inducing birth defect phenotypes.

### Placental crossing and D and X category predictions for small molecules

Using unsupervised learning we generated placental crossing scores and D and X category scores for all FDA-approved and preclinical compounds profiled by LINCS that are included in the ReproTox KG. To obtain true positives for placental crossing, we first extracted the list of 248 compounds assembled by Di Filippo et al. [45]. Category D and X drugs were obtained from DrugCentral [23] and Drugs.com [46], and drugs were filtered by those which could be mapped to the LINCS compounds. Drugs associated with both categories were considered category X. Predictions were made with a symmetric drug-drug similarity matrix using the same approach described by DrugShot [25] by selecting the vectors corresponding to drugs known to be in the true positive set and computing the average similarity score to all drugs in the matrix. The diagonal is set to zero to prevent contribution from the drug itself. Using the similarity scores, receiver operating characteristic (ROC) curves were computed along with the area under that curve (ROC-AUC). Additionally, the percentage of hits in the top 1% were computed by sorting the true scores and computing the sum across the first 1% entries in this vector. This approach was used to score all LINCS compounds. The placental crossing scores and the category D and X scores for all drugs and small molecules in the ReproTox KG are displayed as node properties for drugs and are depicted as the hue level of the drug nodes in the ReproTox KG user interface. In addition, all the predictions are provided as downloadable files available from https://maayanlab.cloud/reprotox-kg/downloads.

### Combining predictions from expression similarity and structural similarity

Two methods were developed to combine the predictions made by the gene expression and chemical structure similarity predictions. Given two scoring vectors produced by the two different similarity matrices, the Top Rank method takes the highest ranking of the drugs across all predictions to be the aggregated score. This score is then used for the ROC curve and ROC-AUC calculation. Alternatively, given two similarity score vectors, one based on expression, and one based on structure, we aggregated these predictions by assigning a weight to each score coming from the two sources: expression and structure. These weights were optimized for performance using the Adam optimizer [47]. The learned weights are then applied for combining the L1000 and structural features in the FDA drug categorization and placenta crossing sets into a singular score. This score is then used for the ROC curve and ROC-AUC calculation.

### UMAP visualization of L1000 perturbations

Uniform Manifold Approximation and Projection (UMAP) [48] was applied to the normalized L1000 count matrix of over 718,055 chemical perturbations performed with different drugs across different cell lines, time points, and concentrations. Perturbations with FDA drug categories D and X and drugs known to cross the placenta were colored by category. To identify the top MOAs in the L1000 perturbation space, we first clustered L1000 perturbations directly using HDBSCAN [49] with a minimum cluster size of 40, we then selected the top 25 clusters with highest concentration of drugs for each drug category, finally we identified the top 5 MOAs for drugs in those clusters. We colored the L1000 UMAP with those top MOAs.

### ReproTox KG backend KG database

The ReproTox KG uses a graph-structured data model to integrate data. The KG is implemented using the Neo4J [50]. The information in the ReproTox KG represents a network of nodes representing birth defects, genes, and drugs, and edges representing their relationships. In addition, attributes/properties of the nodes and edges are provided. The ReproTox KG is made up of datasets from the various sources listed above and listed in two tables (Table 1 and Table 2) and illustrated in the associated schematic (Fig. 1). The ReproTox KG uses standardized JSON schema serialization to ingest data into the KG. Queries to the Neo4J platform are constructed using the Cypher query language [51].

**Table 1.**
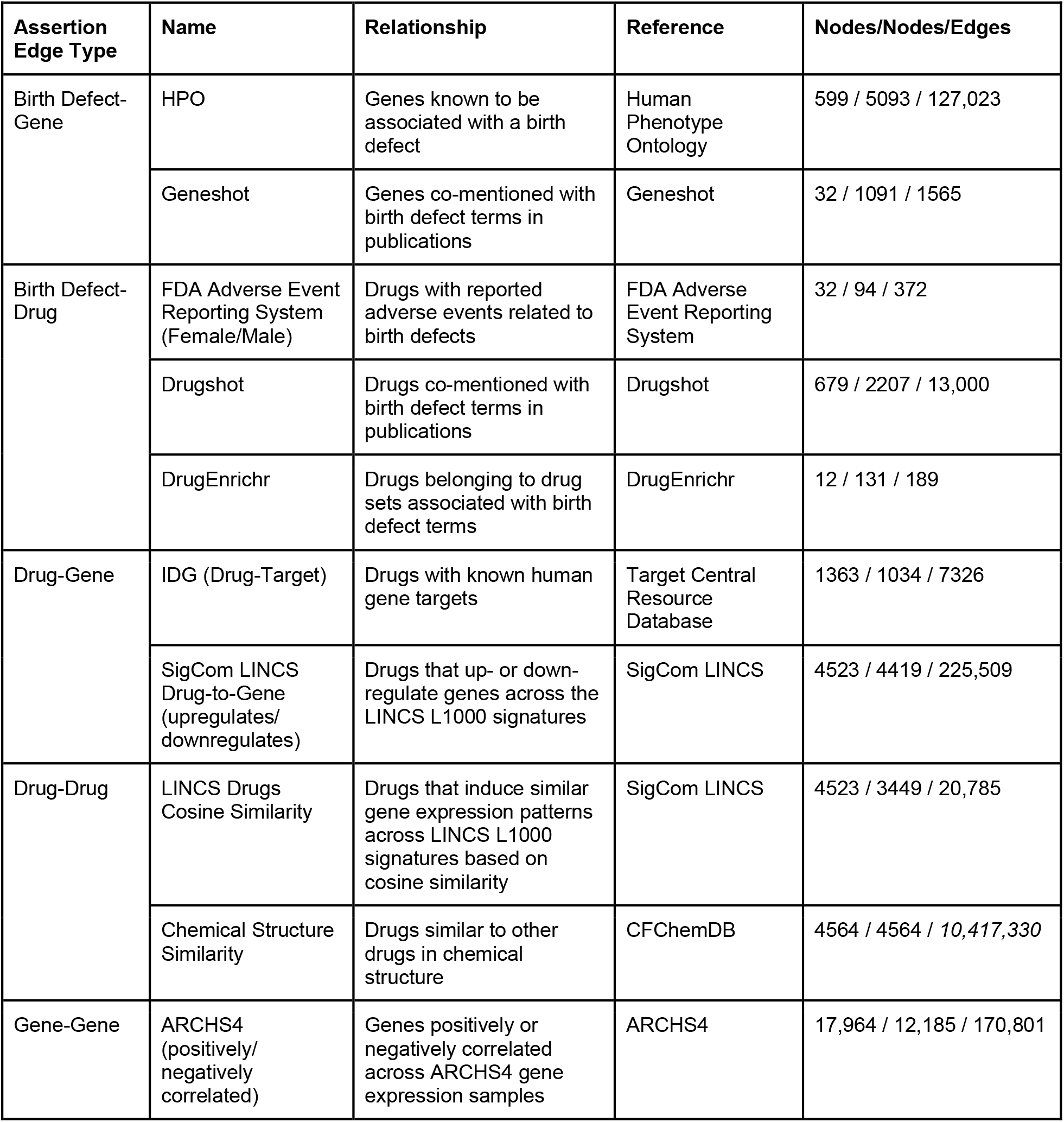
The ReproTox KG is made of entities (nodes) representing birth defects, genes, and drugs that are connected based on semantic assertions (edges/relationships) extracted from different sources. The table lists the type of assertion, the nature of the relationship, the original source from where the assertion was extracted from, and the number of entities and relations for each entry in the table (left entity/right entity/relations).

**Table 2.** The ReproTox KG is made of entities (nodes) representing birth defects, genes, and drugs that are decorated with attributes and properties associated with them, for example, common identifiers. The table lists the properties and their sources for each entity type, namely, birth defects, genes, and drugs represented in the ReproTox KG.

| Entity Type | Property | Source | Description | Entities with property |
| --- | --- | --- | --- | --- |
| Birth Defect | MEDDRA code |  | MedDRA ontology identifier | 32 |
| Drug | Placenta crossing likelihood score | Drugshot | Cosine similarity score to L1000 gene expression signatures for drugs known to cross placental barrier | 4523 |
|  | Placenta crossing likelihood rank | Drugshot | Rank (1=most similar) of cosine similarity score to L1000 gene expression signatures for drugs known to cross placental barrier | 4523 |
|  | SMILES | PubChem | SMILES structure |  |
| Gene | pLI | Exome Aggregation Consortium (ExAC) | Probability of loss of function intolerance | 9466 |
|  | pHI | Collins et al. 2021 [43] | Haploinsufficiency score | 9466 |
|  | pTS | Collins et al. 2021 [43] | Triplosensitivity score | 9466 |
|  | Residual Variant Intolerance Score | NHLBI Exome Sequencing Project | Ratio of functional mutations to total variant sites | 9466 |
|  | Residual Variant Intolerance Score Percentile | NHLBI Exome Sequencing Project | Percentile of RVIS across all scored genes | 9466 |

**Fig. 1.**
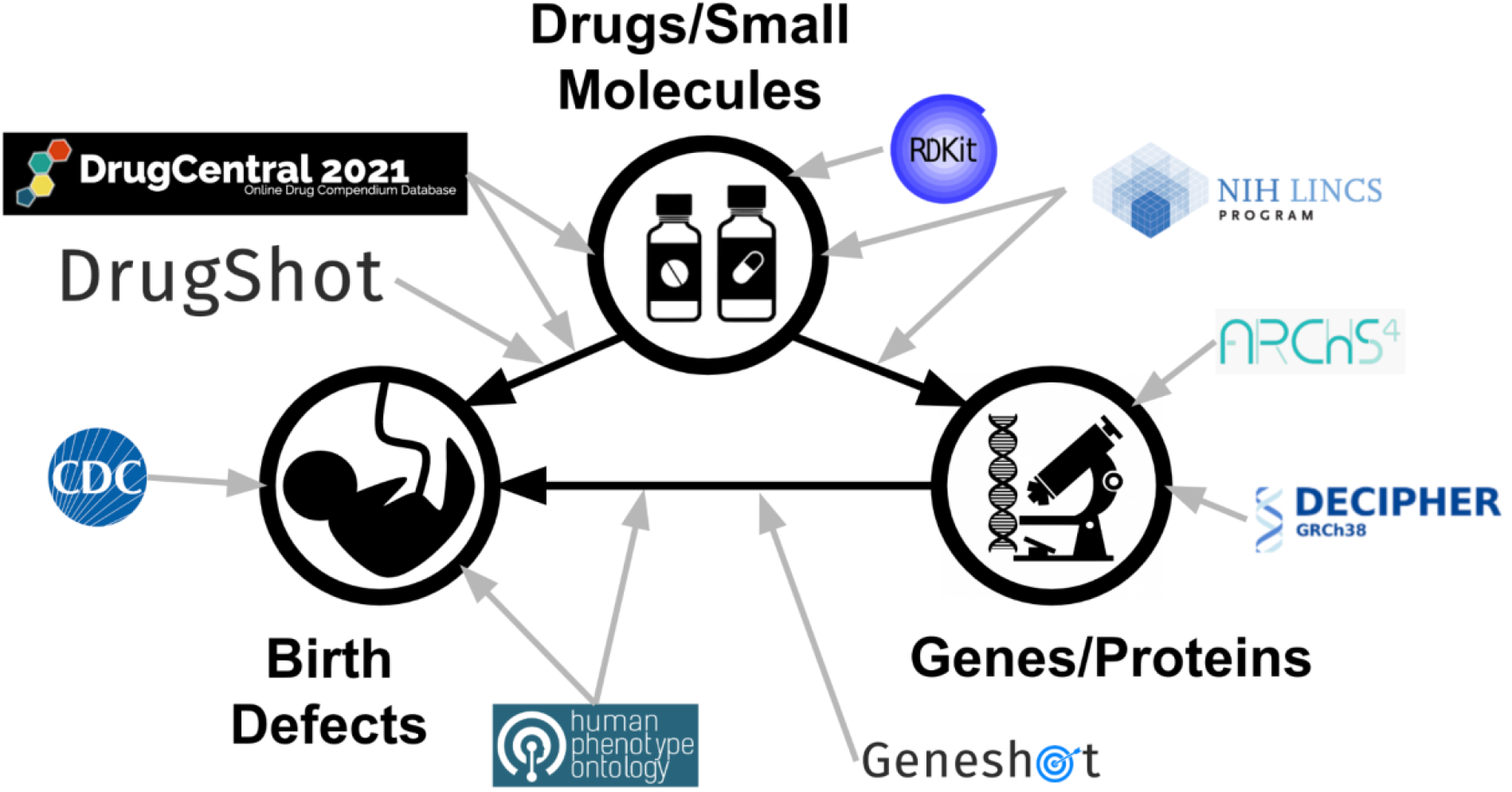
The ReproTox KG is made of lists of birth defects extracted from HPO and the CDC; birth defect gene associations from HPO and Geneshot; drug-birth defect associations from DrugCentral and DrugShot; drug-gene associations from LINCS L1000 data and from drug-target knowledge.

### Original graphical user interface to interact with the ReproTox KG

Since Neo4j currently does not provide an open-source, free, and customizable standalone web-based user interface (UI) to visualize the results from Cypher queries, we developed an original UI with these features for this project. Leveraging the Cytoscape.JS library [52], the UI renders Cypher query results in JSON format into nodes and links network visualizations. The UI provides access to perform queries for finding neighbors of single entities, finding shortest paths between pairs of entities, displaying the networks using various layouts, expanding, and shrinking the size of the displayed subnetwork, viewing properties of nodes and links, and downloading the displayed associations in tabular format.

## Results

### Overall construction and composition of the ReproTox KG

The ReproTox KG contains semantic assertions that connect birth defects, genes, and drugs. In addition, drug-drug and gene-gene similarity assertions are included (Fig. 1; Tables 1-2). Each entity in the ReproTox KG has a set of attributes and properties. Some of these attributes are unique to the project. For example, we ranked the likelihood of all included compounds and drugs to cross the placental barrier and to cause birth defects using an unsupervised machine learning approach. To achieve this, we first identified a list of 248 drugs that are known to cross the placenta [45], and lists of FDA approved drugs classified in the X (n=60) and D categories (n=112). We then mapped these drugs to all the drugs and small molecules profiled by the LINCS L1000 assay (Fig. 2). Next, we constructed two drug-drug similarity matrices, one based on drug structural similarity, and one based on gene expression induced signature similarity. These matrices were used to perform unsupervised machine learning to prioritize all drugs for the likelihood to cross the placenta, or to be categorized as D and/or X. Before performing such predictions with these two matrices, we projected the known placental crossing drugs (Fig. 3A) and the category D and X drugs (Fig. 3B) onto the LINCS L1000 gene expression space of 718,055 gene expression signatures induced by >30,000 small molecules using UMAP [48]. We observe that these drugs fall into distinct regions within the L1000 gene expression space. By comparing the UMAP visualization of the known placental crossing drugs and the category D and X drugs to the same layout with highlighted known mechanisms of actions (Fig. 3C), we observe that dense clusters of D and X drugs involve estrogen disruptors and topoisomerase inhibitors. Other clusters colored by their unique MOAs also have many placental crossing drugs and category D and X drugs within them. The observed punctate distribution strongly suggests that we can make predictions about the likelihood of preclinical drugs to induce birth defects and cross the placenta.

**Fig. 2.**
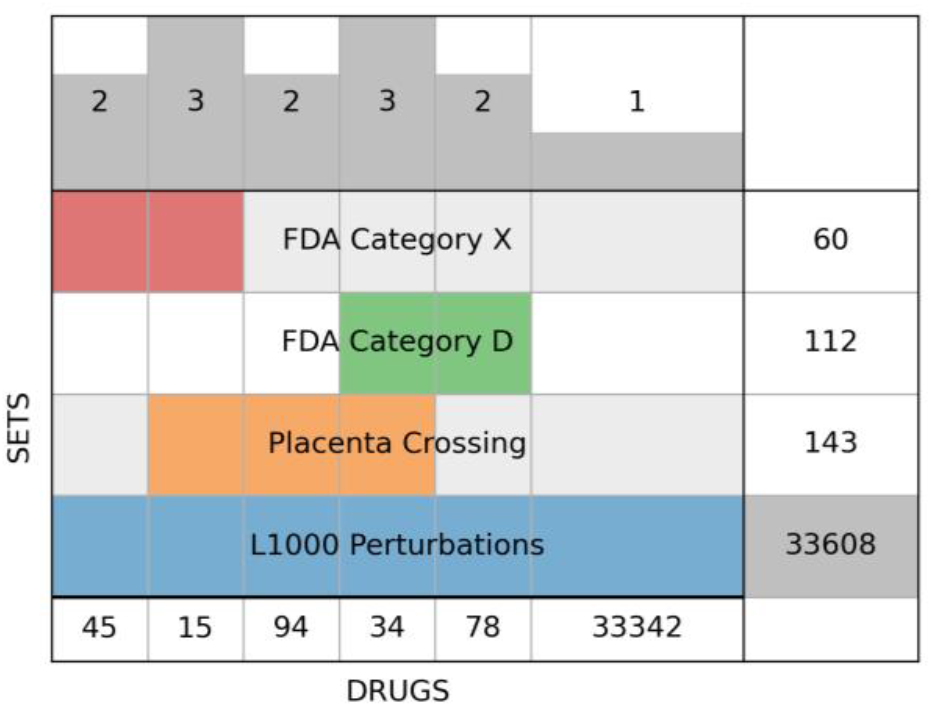
Supervenn diagram of drug identifier overlap between FDA category D and X, known placenta crossing drugs, and unique drugs and small molecules within the L1000 LINCS perturbation datasets. Drugs and compounds not represented in the L1000 perturbations are not shown or considered.

**Fig. 3.**
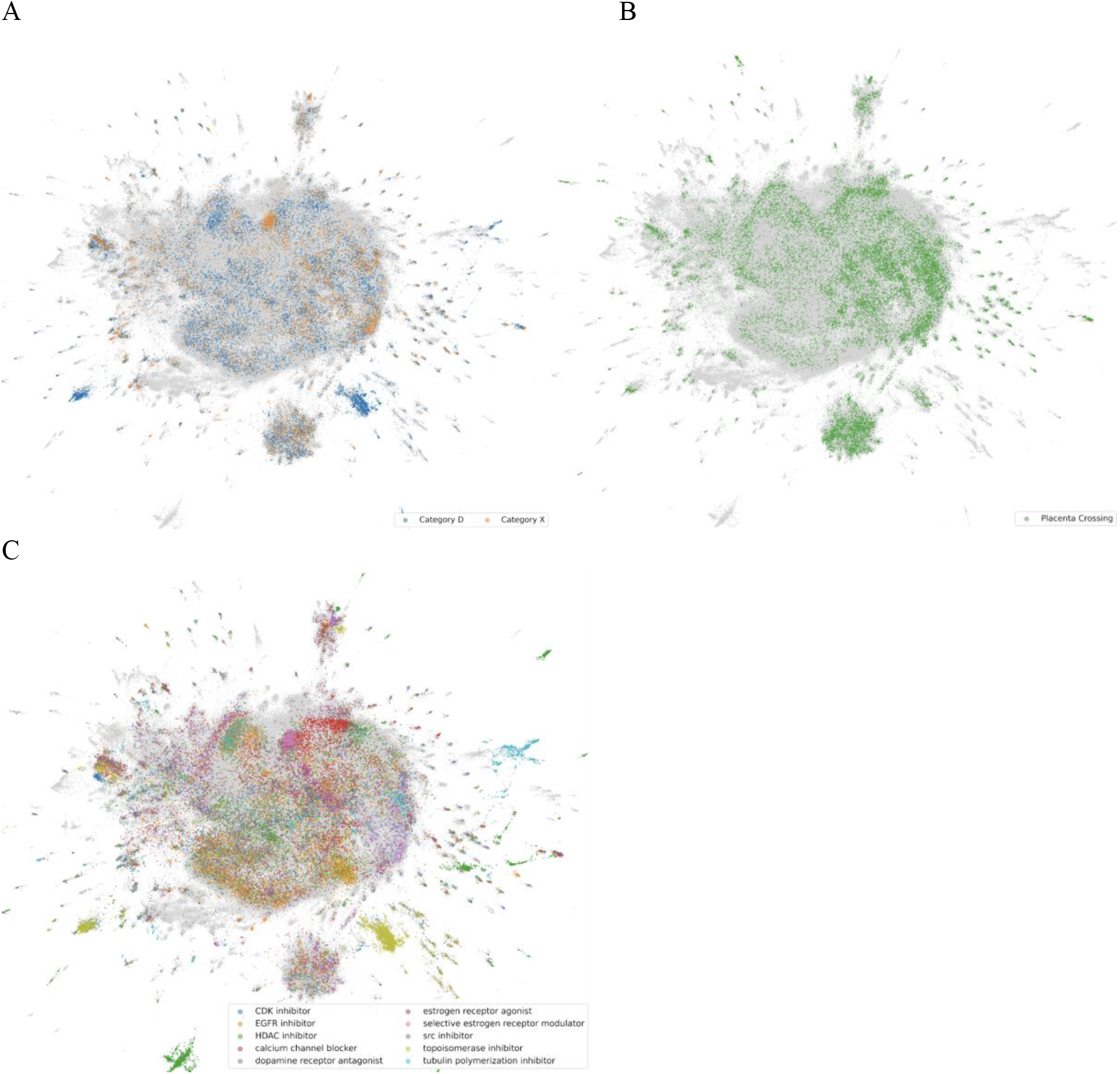
UMAP of 718,055 L1000 perturbations, colored by (A) FDA D and X category; (B) known placental crossing; (C) top MOAs across clusters. Clusters computed using HDBSCAN with a minimum cluster size of 40, top 25 clusters for each category and top 5 MOAs of those clusters are included.

Next, we applied the unsupervised learning approach to rank all mapped approved drugs and preclinical compounds to predict their likelihood to cross the placental barrier or induce a birth defect. We observed that with the L1000 signature similarity matrix we achieve an AUC of 0.620 for predicting D and X category membership, and 0.725 for placental crossing (Fig. 4A, Fig. 4C). The predictions that are based on chemical structural similarity achieve AUCs of 0.746 and 0.660 for D and X category membership and for placental crossing, respectively (Fig. 4A and 4C). Combining the predictions made by gene expression with chemical structure together with the Top Rank or the weighted contribution methods improved such predictions to 0.803 and 0.788 for D and X category membership, as well as 0.785 and 0.759 for the placental crossing predictions, respectively. Overall, these are high quality predictions for an unsupervised approach. Importantly, these predictions perform well at the leading edge (Fig. 4B, Fig. 4D, and Tables 3 and 4). It should be noted that predictions made with structural similarity only performed well when we defined the similarity between compounds using IDF instead of Tanimoto. This is likely because there is a bias with the Tanimoto method which emphasizes similarity between complex larger compounds that share common features. The predictions made by the unsupervised method highly ranked compounds that are known as ACE inhibitors, antibiotics, and statins (Tables 3 and 4). This is not surprising because such compounds are already common among known category X & D drugs [53] and drugs that are known to cross the placenta. For example, the top ranked drug by structural similarity to be categorized as X & D is enalaprilat. Enalaprilat is a known ACE inhibitor used to treat high blood pressure and is administered via IV [54]. It is listed as category C for the first trimester and as category D for the second and third. Overall, such predictions can be used to warn about the potential of newly approved drugs to cross the placenta and induce birth defects.

**Fig. 4.**
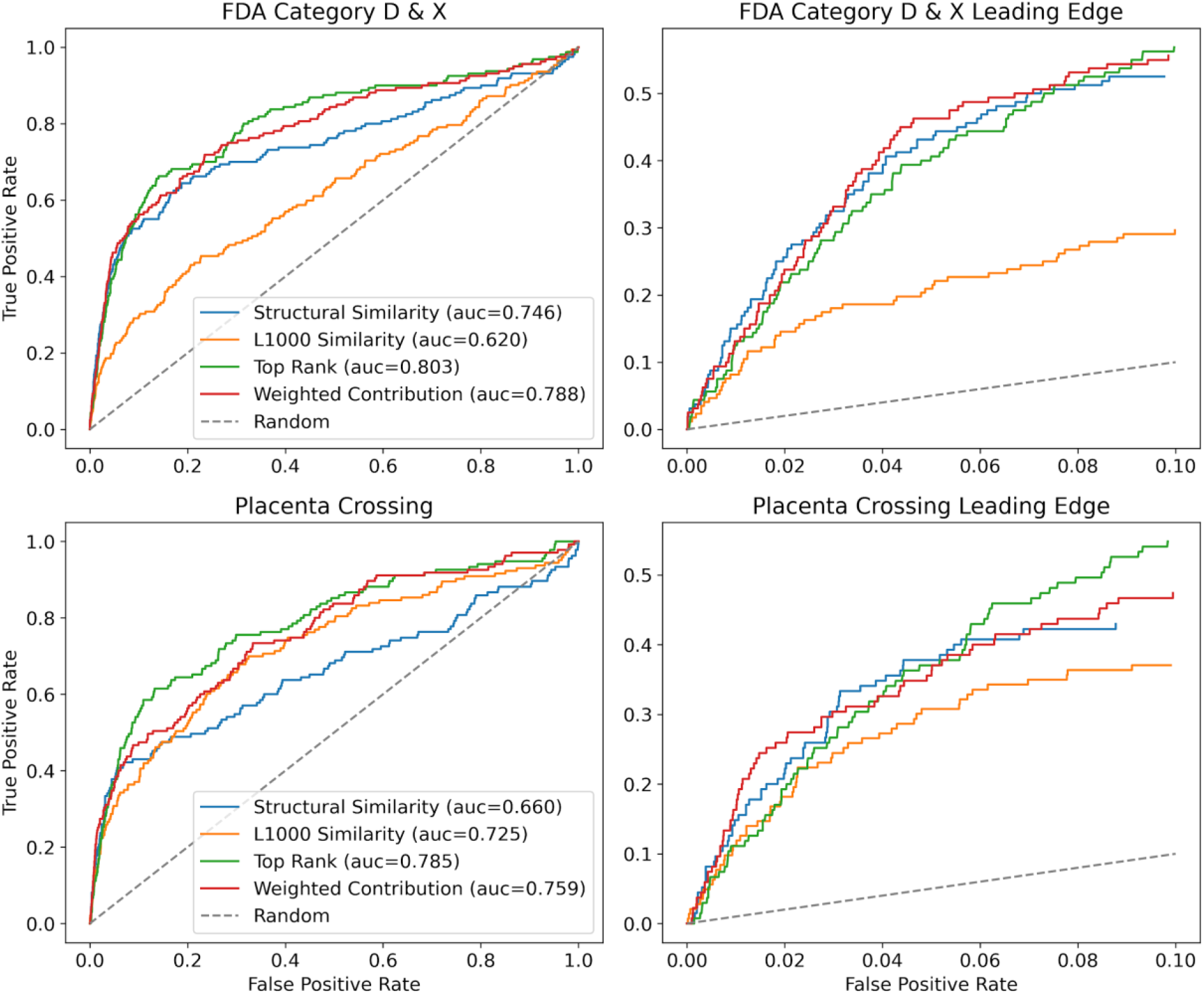
ROC curves colored by prediction method for predicting FDA D and X categories (top) and placenta crossing (bottom). AUC values shown in the legend. Leading edges of the same ROC curves are shown on the right of each complete ROC plot.

**Table 3.** Top predicted X and D category drugs and preclinical compounds. The top 15 ranked compounds predicted using unsupervised learning with L1000 gene expression similarity or chemical structure similarity, or two by two methods that combine the predictions from the two sources, namely, weighted, and top rank, are listed together with whether these were previously known to belong to the X or D categories.

| Drug | D & X<br>L1000 | D & X<br>Structure | Weighted | Top Rank | D Known | X Known |
| --- | --- | --- | --- | --- | --- | --- |
| enalaprilat |  | 1 | 1 | 1 | 0 | 0 |
| TAK-715 | 1 |  |  | 1 | 0 | 0 |
| pitavastatin |  | 2 | 11 | 2 | 0 | 0 |
| BRD-K08703257 | 2 |  |  | 2 | 0 | 0 |
| ramipril |  | 3 | 2 | 3 | 1 | 0 |
| pentoxifylline | 3 |  |  | 3 | 0 | 0 |
| trandolapril |  | 3 | 6 | 4 | 1 | 0 |
| troglitazone | 4 |  |  | 4 | 0 | 0 |
| perindopril |  | 4 | 3 | 5 | 1 | 0 |
| FTI-276 | 5 |  |  | 5 | 0 | 0 |
| BRD-K76846644 |  | 5 | 5 | 6 | 0 | 0 |
| Gossypetin | 7 |  |  | 6 | 0 | 0 |
| pravastatin |  | 6 | 4 | 7 | 0 | 1 |
| phenothiazine | 9 |  |  | 7 | 0 | 0 |
| evodiamine |  | 7 | 7 | 8 | 0 | 0 |
| lorazepam | 10 |  |  | 8 | 1 | 0 |
| BRD-K40167599 |  | 8 |  | 9 | 0 | 0 |
| losartan | 11 |  |  | 9 | 1 | 0 |
| lovastatin |  | 9 |  | 10 | 0 | 1 |
| zebularine | 13 |  |  | 10 | 0 | 0 |
| CTPB |  | 10 | 15 | 11 | 0 | 0 |
| risperidone |  |  |  | 11 | 0 | 0 |
| VX-765 |  | 11 | 8 | 12 | 0 | 0 |
| BMS-191011 |  |  |  | 12 | 0 | 0 |
| methiopril |  | 12 | 10 | 13 | 0 | 0 |
| EX-527 |  |  |  | 13 | 0 | 0 |
| canrenone |  | 13 | 12 | 14 | 0 | 0 |
| BRD-K55591206 |  |  |  | 14 | 0 | 0 |
| BRD-K80672993 |  |  |  | 15 | 0 | 0 |
| oxytetracycline |  | 14 |  | 15 | 0 | 0 |
| BRD-K63784565 |  |  | 9 |  | 0 | 0 |
| zofenopril-calcium |  |  | 13 |  | 0 | 0 |
| BRD-K93367411 |  |  | 14 |  | 0 | 0 |
| BRD-K08533345 | 14 |  |  |  | 0 | 0 |
| BRD-K27194553 | 12 |  |  |  | 0 | 0 |
| BRD-K27889081 | 8 |  |  |  | 0 | 0 |
| BRD-K37848780 | 15 |  |  |  | 0 | 0 |
| BRD-K76551930 | 6 |  |  |  | 0 | 0 |
| imidapril |  | 15 |  |  | 0 | 0 |

**Table 4.** Top predicted placental crossing drugs and preclinical compounds. The top 15 ranked compounds predicted using unsupervised learning with L1000 gene expression similarity or chemical structure similarity, or two by two methods that combine the predictions from the two sources, namely, weighted, and top rank, are listed together with whether these were previously known to cross the placenta.

| Drug | L1000 | Structure | Weighted | Top Rank | Known |
| --- | --- | --- | --- | --- | --- |
| nafcillin |  | 1 | 1 | 1 | 0 |
| FTI-276 | 1 |  |  | 1 | 0 |
| piperacillin |  | 2 | 4 | 2 | 0 |
| Gossypetin | 2 |  |  | 2 | 0 |
| cefotaxime |  | 3 | 3 | 3 | 0 |
| TAK-715 | 3 |  |  | 3 | 0 |
| ciclacillin |  | 4 | 2 | 4 | 0 |
| BRD-K08703257 | 4 |  |  | 4 | 0 |
| 7-aminocephalosporanic-acid |  | 5 | 6 | 5 | 0 |
| temozolomide | 5 |  |  | 5 | 0 |
| penicillin |  | 6 | 5 | 6 | 0 |
| CGS-21680 | 6 |  |  | 6 | 0 |
| ceforanide |  | 7 | 9 | 7 | 0 |
| Y-27632 | 7 |  |  | 7 | 0 |
| lorazepam | 8 |  |  | 8 | 1 |
| BRD-K43966364 |  | 8 | 8 | 9 | 0 |
| benzathine |  | 8 |  | 9 | 0 |
| estradiol-cypionate | 9 |  |  | 10 | 0 |
| BRD-K50776152 | 10 |  |  | 11 | 0 |
| isoetharine |  | 9 |  | 11 | 0 |
| enalaprilat |  | 10 | 12 | 12 | 0 |
| rolipram | 11 |  |  | 12 | 0 |
| BRD-A66025870 |  | 11 |  | 13 | 0 |
| PT-630 | 12 |  |  | 13 | 0 |
| EMF-csc-9 | 13 |  |  | 14 | 0 |
| practolol |  | 12 |  | 14 | 0 |
| cephalothin |  | 13 | 15 | 15 | 1 |
| DL-TBOA | 14 |  |  | 15 | 0 |
| pravastatin |  | 14 | 7 |  | 0 |
| cefoperazone |  |  | 10 |  | 1 |
| orciprenaline |  |  | 11 |  | 0 |
| ampicillin |  |  | 13 |  | 1 |
| dicloxacillin |  |  | 14 |  | 0 |
| BRD-K55591206 | 15 |  |  |  | 0 |
| micropenin |  | 15 |  |  | 0 |

In addition, for each gene included in the KG, we computed likelihood for deleterious mutations using three established methods: (pLI) scores [39], RVIS [41], and Dosage sensitivity scores [20]. Each entity in the ReproTox KG also includes links out to databases based on entity ID resolution. In particular, 694 birth defects are mapped to HPO identifiers [18], 18,233 genes and proteins are mapped to HGNC IDs, and 5,403 drugs are mapped to their PubChem identifiers [34]. Lists of birth defect terms were extracted from HPO [18] and the CDC website [19]. The 127,023 associations between birth defect terms and genes were extracted from OMIM [26], Orphanet [27], ClinVar [28], DISEASES [23], DECIPHER [30], American Heart Association (AHA) [31] and Geneshot [32]. The 13,561 assertions between birth defects and drugs were extracted from DrugCentral [32], DrugShot [25], DrugEnrichr, and FAERS [24]. Two types of assertions connect genes and drugs within the ReproTox KG, genes that are differentially expressed after drug treatment based on transcriptomics and known drug targets for the drugs. Overall, 225,509 drug-gene associations were extracted from the LINCS L1000 data [36], and 7,326 drug-target assertions were extracted from TCRD [16]. Similarly, 9,546 drug-drug similarity assertions were identified based on chemical similarity and 33,608 based on gene expression signature similarity. Finally, gene-gene similarity included in the KG is based on gene-gene co-expression [44].

The processed data from these resources was created by customized extract, transform, and load (ETL) scripts and stored in a JSON schema data model. This processed data was ingested into a Neo4J database, and it is made available for download on the ReproTox KG website at: https://maayanlab.cloud/reprotox-kg/downloads. The ETL scripts are open source and available from: https://github.com/nih-cfde/ReproToxTables/. To provide access to the processed data in a user-friendly manner, we developed an original graphical user interface (Fig. 5).

**Fig. 5.**
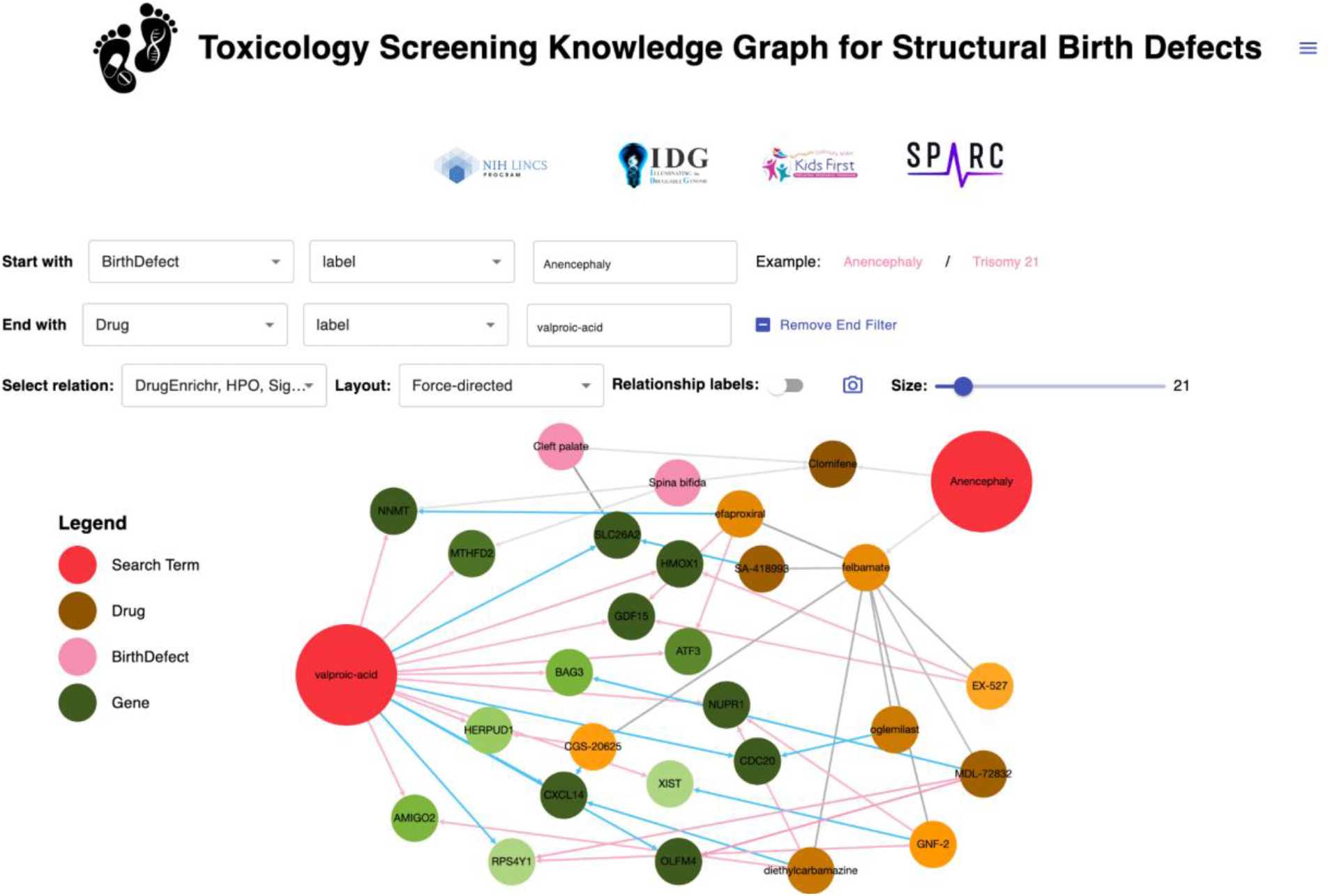
Screenshot from the ReproTox KG user interface. A query to identify connections between the birth defect anencephaly and the drug valproic acid with a limit of 21 nodes is provided as an example.

### Extraction of birth defect-gene-small molecule cliques to explain potential MOAs

To demonstrate the utility of the ReproTox KG to illuminate new knowledge, we queried the graph to identify all three-node cliques/cycles. That is, we extracted from the ReproTox KG all instances where a birth defect was connected to a gene and a drug that were also connected. In total, 533 such cliques/cycles were identified. From this collection of cliques/cycles, we identified six drugs and small molecules that were not previously listed as crossing the placenta and had a placental scoring rank in the top 10% (<3000 out of ∼30,000) (Fig. 6). This subnetwork demonstrates how the ReproTox KG can be used to suggest mechanisms of action (MOA) for how drugs and pre-clinical compounds may induce specific birth defects by affecting the gene expression of genes already known to be associated with the birth defect. For example, LINCS L1000 transcriptomics data shows that the approved drug methotrexate, a chemotherapeutic and immunosuppressive drug, inhibits the expression of the mitotic checkpoint serine/threonine-protein kinase BUB1. BUB1 is known to cause microcephaly when mutated [55], and methotrexate is known to cause microcephaly and atrial defects [56]. Hence, this adverse effect of methotrexate can be attributed to its direct influence on the expression levels of BUB1. Similarly, the experimental drug LY-294002 which is a morpholine-containing chemical compound that is a strong inhibitor of PI3K, was previously shown to influence cell proliferation of epithelial cells isolated from human fetal palatal shelves (hFPECs) [57]. Besides inhibiting the activity of PI3K, LY-294002 increases the expression of DUSP6, a dual specificity phosphatase that dephosphorylates members of the PI3K pathway. The approved antidepressant drug sertraline was reported to induce cardiac and vascular birth defects based on analysis of FAERS [58]. The ReproTox KG subnetwork of cliques/cycles suggests that such adverse birth defects could be mediated via the activation of the dehydrocholesterol reductase DHCR7 and DHCR24. Mutations in DHCR7 are known to cause Smith-Lemli-Opitz syndrome, a disease of multiple congenital abnormalities [59], while mutations in DHCR24 can cause desmosterolosis [60]. Hence, it is plausible that sertraline mediates induction of cardiac and vascular birth defects via its up-regulatory effects on DHCR7 and DHCR24. Overall, these are just a few examples of how the ReproTox KG can illuminate new knowledge about potential mechanisms of how drugs and preclinical small molecules may induce birth defects.

**Fig. 6.**
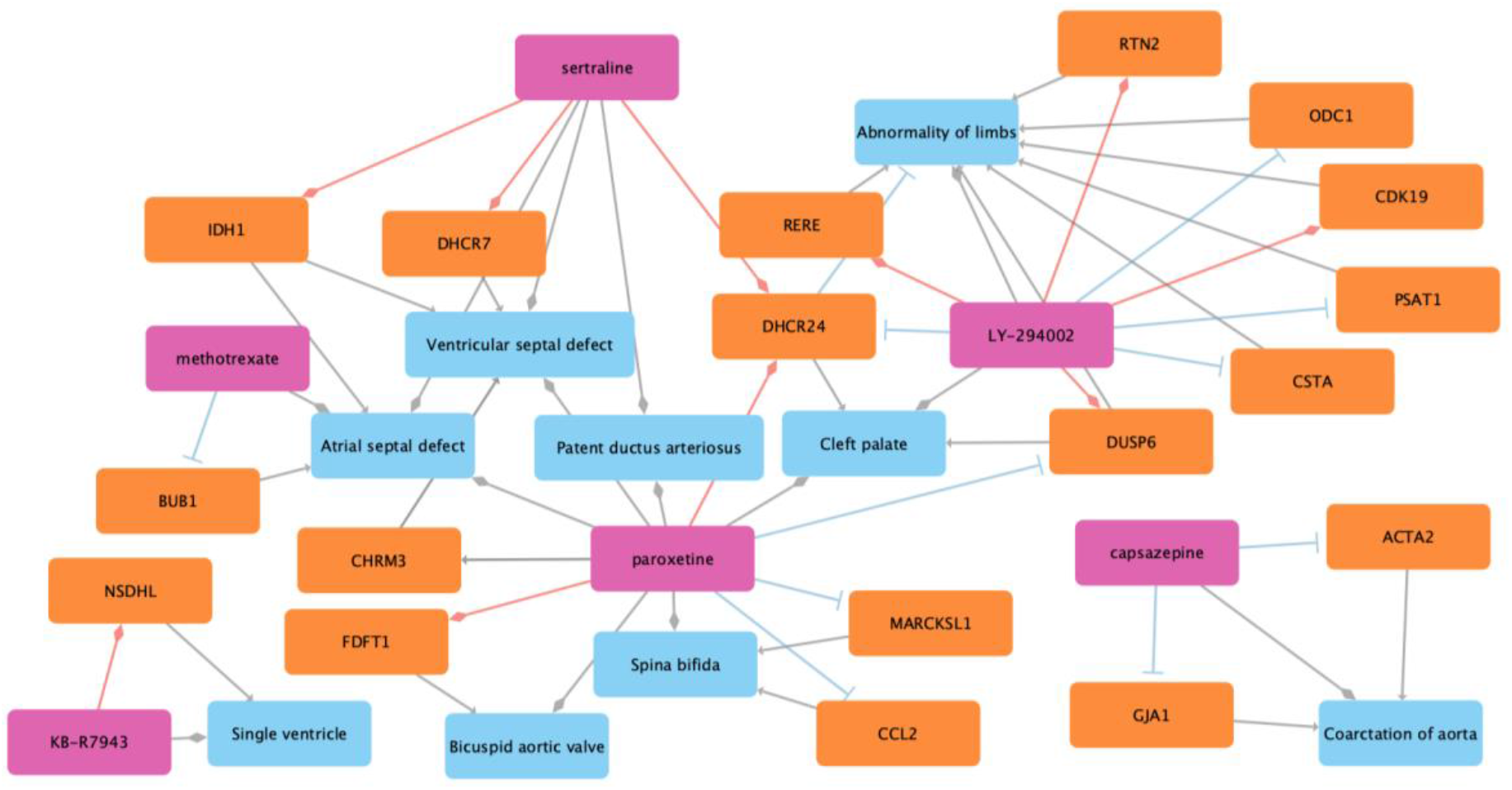
Cliques of drugs with a placenta crossing predicted ranks of less than 3000, that are also known to induce a birth defect based on literature evidence, connected to the genes that their expression is affected based on LINCS L1000 data, and associations between those genes where known mutations are also associated with the same birth defect. Light blue nodes represent birth defect terms, orange nodes represent genes, and pink nodes represent drugs and preclinical small molecules. Red lines with diamond arrowheads indicate an L1000 consensus drug signature that up-regulates the gene, and plungers indicate an L1000 consensus drug signature that down-regulates the target gene. Gray arrowheads indicate genes that their mutations induce a birth defect, and gray diamond-heads connect drugs to the birth defects they are known to induce.

## Discussion and Conclusions

To characterize associations between small molecule compounds and their potential to induce reproductive toxicity, we gathered knowledge from multiple sources to construct a reproductive toxicity Knowledge Graph with an initial focus on associations between birth defects, drugs, and genes. The idea of abstracting genes, drugs, and diseases into networks is not new. We and others constructed networks to represent functional and physical associations between genes/proteins [61] [62] [63], drugs and their targets [64] [65], and diseases based on their gene set similarity [66]. The unique features of the ReproTox KG are that it provides a flexible framework not only to connect entities such as gene-drug, gene-gene, gene-birth defect, drug-drug, and drug-birth defects, but also a flexible way to query this network, extend it, visualize it, and add attributes to different node and link types. The ReproTox-KG is an initial effort towards integrating knowledge about birth defects, genes, and drugs. Similar efforts have been recently published, including studies that attempted to use graph embedding algorithms to predict missing/novel associations between drugs and diseases [67], for drug repurposing opportunities [68] [69], predicting drug targets [70] [71], adverse events [72], and drug-drug interactions [73]. These are just a few studies in this domain. Here, we did not attempt to make predictions based on the knowledge graph structure but provided the needed building blocks to enable such future applications. Hence, the ReproTox KG was developed as a resource for the community to explore and expand.

One of the limitations of ReproTox KG, and knowledge graphs representation in general, is the ability to cover many associations between entities. For example, we decided to only consider the top 25 up- and down-regulated genes for each drug. This leaves out many genes that may be affected by drugs but will be missed from queries and post-hoc analyses. We also created consensus signatures for each drug from the LINCS L1000 data, this approach masks the effect of drugs in specific cellular contexts. This was done to make the ReproTox KG project focused and manageable. However, tissue and cell type distribution of the affected genes, and how drugs and small molecules induce such differential effects, are critical information for associating genes and drugs with birth defects. Such information is partially available and could be included in future releases of ReproTox KG. One excellent resource for gene expression during development is DESCARTES, a human cell atlas for fetal tissues [74].

In conclusion, ReproTox KG provides a resource for exploring knowledge about the molecular mechanisms of birth defects with the potential of predicting the likelihood of genes and preclinical small molecules to induce birth defect phenotypes. It should be noted that the ReproTox KG is preliminary and should not be used for clinical applications and clinical decision support.

## Acknowledgements

This project was supported by NIH grants OT2OD030160, OT2OD030546, OT2OD032619, and OT2OD030162.

